# Can we harness digital technologies and physiology to hasten genetic gain in U.S. maize breeding?

**DOI:** 10.1101/2021.02.23.432477

**Authors:** C. Diepenbrock, T. Tang, M. Jines, F. Technow, S. Lira, D. Podlich, M. Cooper, C. Messina

## Abstract

Genetic gain in breeding programs depends on the predictive skill of genotype-to-phenotype algorithms and precision of phenotyping, both integrated with well-defined breeding objectives for a target population of environments (TPE). The integration of physiology and genomics could improve predictive skill by capturing additive and non-additive interaction effects of genotype (G), environment (E), and management (M). Precision phenotyping at managed stress environments (MSEs) can elicit physiological expression of processes that differentiate germplasm for performance in target environments, thus enabling algorithm training. Gap analysis methodology enables design of GxM technologies for target environments by assessing the difference between current and attainable yields within physiological limits. Harnessing digital technologies such as crop growth model-whole genome prediction (CGM-WGP) and gap analysis, and MSEs, can hasten genetic gain by improving predictive skill and definition of breeding goals in the U.S. maize production TPE. A half-diallel maize experiment resulting from crossing 9 elite maize inbreds was conducted at 17 locations in the TPE and 6 locations at MSEs between 2017 and 2019. Analyses over 35 families represented by 2367 hybrids demonstrated that CGM-WGP offered a predictive advantage (*y*) compared to WGP that increased with occurrence of drought as measured by decreasing whole-season evapotranspiration (ET; log(*y*) = 0.80(±0.6) − 0.006(±0.001) × *ET*; *r*^2^ = 0.59; *df* = 21). Predictions of unobserved physiological traits using the CGM, akin to digital phenotyping, were stable. This understanding of germplasm response to ET enables predictive design of opportunities to close productivity gaps. We conclude that enabling physiology through digital methods can hasten genetic gain by improving predictive skill and defining breeding objectives bounded by physiological realities.

## Introduction

The combination of molecular technologies and digital prediction methodologies has transformed crop improvement over the last decade (Cooper et al., 2014b; Poland, 2015; Ramirez-Villegas et al., 2020) and increasingly enabled farmers to produce enough food, feed, fuel and fiber for society. However, future agriculture is unlikely to balance supply and demand for food (Ray et al., 2013; Fisher et al., 2014), even in the absence of any considerations to reduce greenhouse gas emissions (NASEM, 2019). Novel systems frameworks are required for society to accelerate genetic gain to deliver on food, nutritional, and economic security as target environments change. Methods to effectively deal with genotype x environment interactions (GxE), which are a major factor limiting realization of the required increases in rate of genetic gain in all major crops (Cooper et al., 1995; Chapman et al., 2000; de la Vega and Chapman, 2001; Cooper et al., 2014a; Mwiinga et al., 2020), have been developed (Heslot et al. 2014; Li et al., 2018; Millet et al., 2019; Monteverde et al. 2019; van Eeuwijk et al., 2019; Robert et al. 2020). However, methods to predict long-term consequences of co-selection of genotypes and optimal agronomic management practices (M), which underpinned the historical high rates of genetic gain for maize yield in the US corn-belt (Duvick, 2005), are only just emerging (Messina et al., 2018; Cooper et al. 2020b). This is a *super wicked* problem because the information needed to train data-driven models is only routinely available for few genotypes and creating training sets for many genotypes could be prohibitively expensive (Fig 1). The integration of physiology-based and data-driven approaches has been proposed as a workable solution whereby scientific understanding can effectively deal with model underdetermination (Messina et al., 2018; Hammer et al., 2019; Messina et al., 2020b; McCormick et al., 2020).

**Fig 1.**
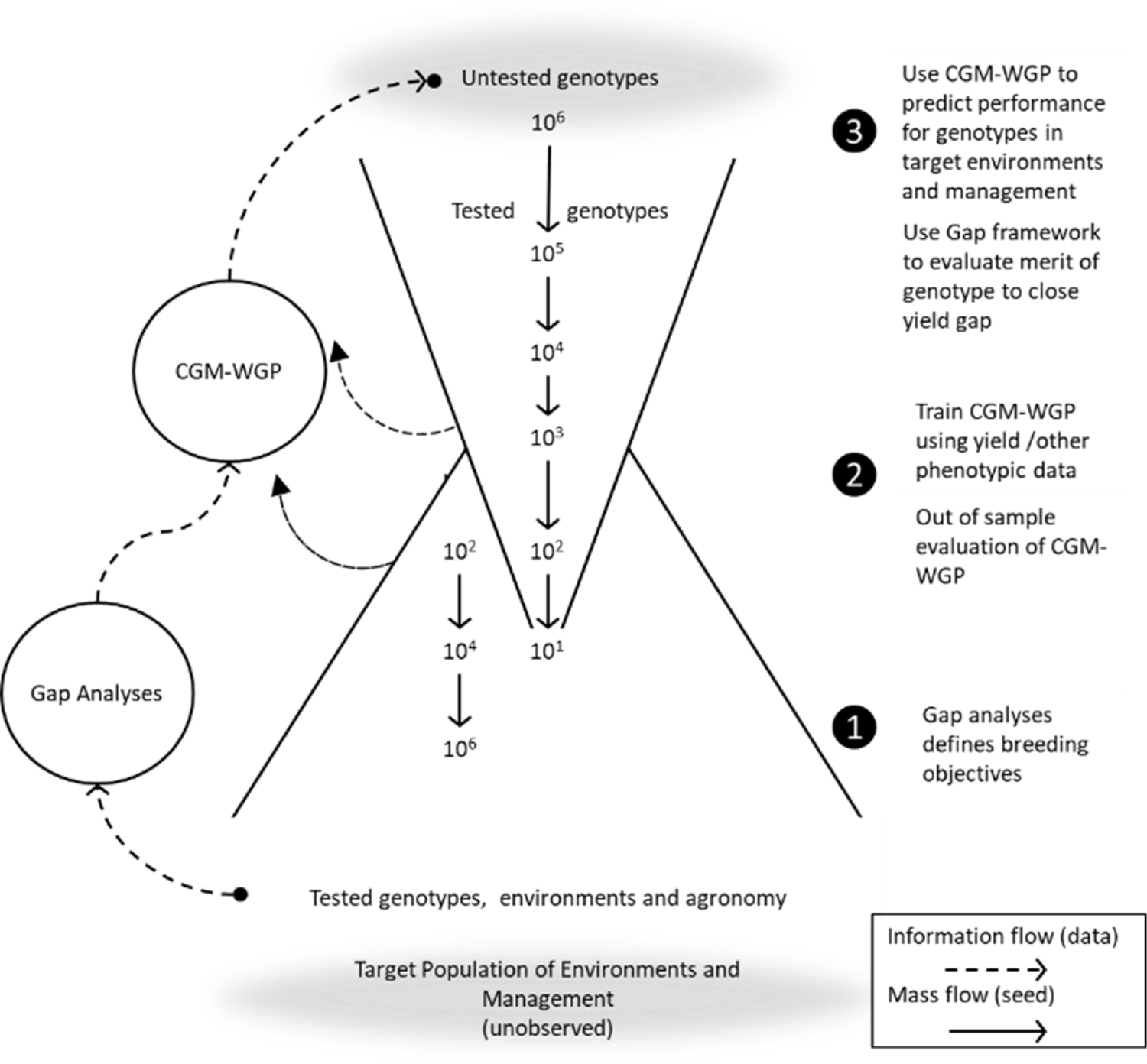
Diagram of order of magnitude for hybrids advancing through product development along with processes to improve breeding efficiencies, Gap analyses and crop growth model – whole-genome prediction (CGM-WGP).

Crop growth models (CGM) are cognitive constructs that capture in mathematical form physiological knowledge with various degrees of detail (Fig 2). Combining a CGM with whole-genome prediction (CGM-WGP) is the extension of the whole-genome prediction (WGP, Meuwissen et al. 2001; Lorenz et al. 2011; Heslot et al. 2012; Poland et al., 2012) framework to integrate plant physiology and genomics encapsulated within crop models. This framework is unique in its conception to leverage fundamental physiological understanding to connect genotypes and phenotypes. Previous demonstration through simulation and empirical studies (Technow et al. 2015, Cooper et al. 2016, Messina et al. 2018), albeit limited in scope, produced encouraging results. These examples used a maize CGM that has been refined over decades of experimentation and contains modules to simulate yield potentials and reductions due to intensity and timing of water stress (Hammer et al., 2009; Messina et al., 2015; Messina et al., 2019). CGM-WGP should be seen as a physiological (Fig. 2a) and quantitative genetics integrated framework (Cooper et al., 2020a; Fig. 2c) that could be useful for germplasm characterization and prediction, with dynamic consideration of the main and interaction effects of G, E and M on crop performance.

**Figure 2.**
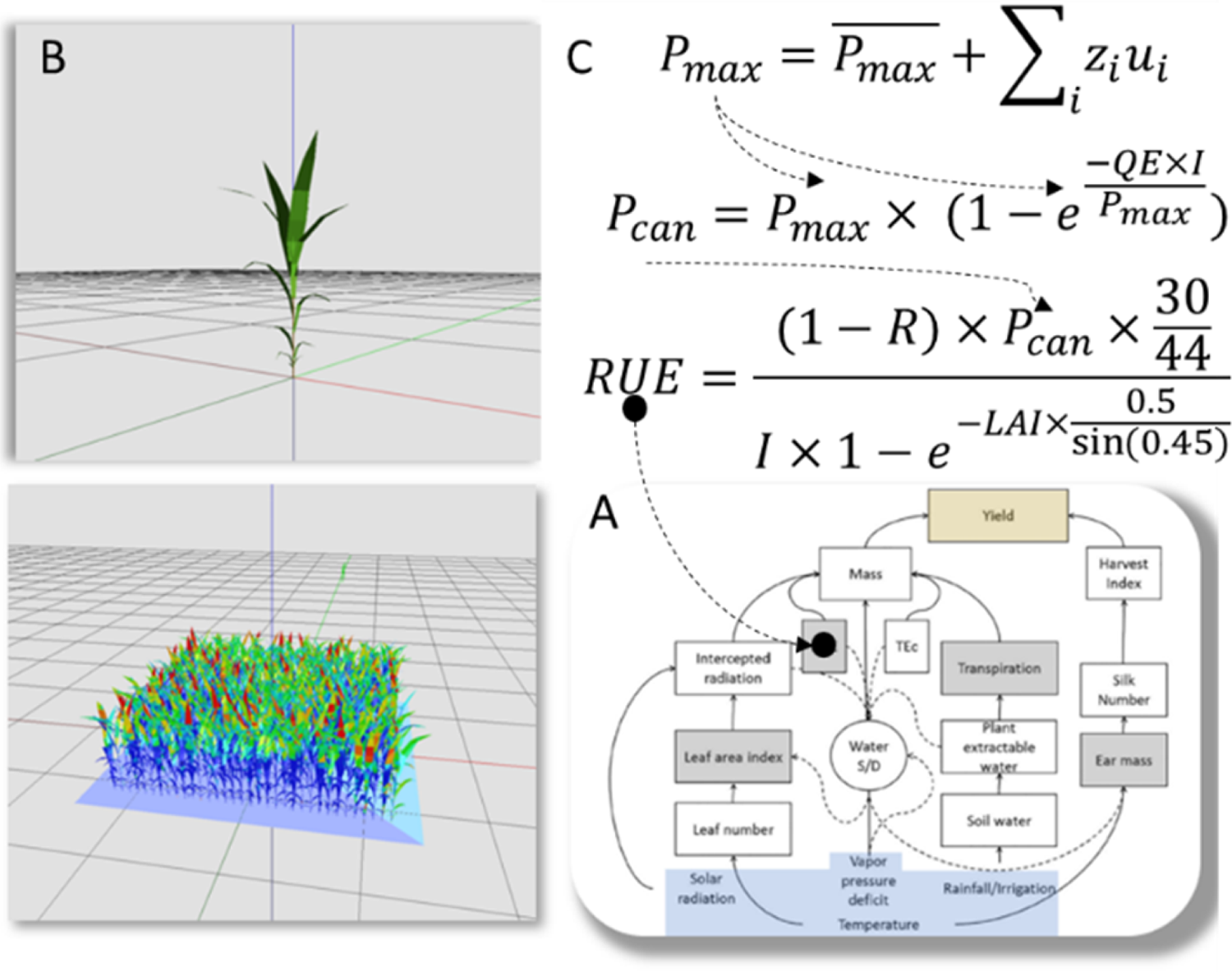
Diagram of a crop growth model (A), leaf emergence and light interception diagrams (B), and genotype◊phenotype connection through an example mapping maximum photosynthesis (*Pmax*), canopy photosynthesis (*Pcan*) and radiation use efficiency (*RUE*). Genotype marker (**z**) and effects (**u**) at genome positions (*i*), *QE*: quantum efficiency, *LAI*: Leaf Area Index, *I*: Light Interception.

In CGM-WGP, through many iterations of CGM runs housed inside of a Metropolis-Hastings-within-Gibbs algorithm, the trait estimates with highest posterior probability for each genotype in a training set (i.e., the set of tested genotypes used in the estimation step) are determined through sampling, for a limited set of physiological traits specified by the practitioner (Messina et al. 2018). These traits should express genetic variation in the target breeding populations and be highly heritable—largely insensitive to the effects of environment and management. Model parameters in the photosynthesis response to CO_2_ and light (Fig. 2) are good examples of processes for which there is reasonable evidence for biophysical and genetic regulation and therefore suitable targets for estimation (Leakey et al., 2006; Wu et al., 2019). Yield predictions can then be generated for field tested or untested individuals, through a final set of runs of the CGM over the distribution of samples obtained in the previous step for each tested or untested environment and agronomic management of interest (Fig 1). In this final prediction step, marker-based estimates (e.g., Fig. 2c) are used to calculate the appropriate value to be used for the physiological trait(s) that were estimated in the training step. Any target environment for which the crop growth model assumptions are appropriate can be run in the prediction step for a defined set of agronomic practices.

Breeders conduct field trials to make inferences regarding the performance of genotypes of interest in certain E and M combinations within the target population of environments (TPE; Comstock, 1977, Cooper and DeLacy 1994). However, even with careful experimental design, any given multienvironment trial (MET) samples a relatively small and often inadequate fraction of the TPE (Cooper et al., 1995; Cooper et al. 2014b; van Eeuwijk et al. 2019). That is, several testing sites that are selected for their potential to represent distinct environment types in the TPE could experience highly similar conditions in a given year, leaving other environment types under-represented (Cooper et al. 2014a). Weighting selection decisions by the frequency of occurrence of environment type was advocated to overcome this problem (Podlich et al., 1999). Other approaches utilize managed stress environments (MSEs) designed to emulate the timing and severity of a stressor (e.g. drought in the flowering stage of development), and/or to elicit a physiological response that separates germplasm for adaptation to the target environmental conditions encountered in the TPE with a high enough frequency (Cooper et al., 1995; Cooper et al., 2014a,b). By generating contrasting environment types through use of MSE management to discriminate germplasm, it is possible to estimate physiological trait values and marker effects that give rise to the manifested norms of reactions characteristic of each genotype for the target environments of the TPE.

The Gap analysis methodology seeks to quantify the difference between realized crop yields and what could be achieved given the availability of limiting natural resources (van Ittersum et al., 2013). This methodology provides an estimate for the realization of both the genetic and environmental potential at any given site and year. Cooper et al. (2020b) proposed to use this framework to design GxM technologies to close productivity gaps; breeding objectives are set relative to potential and realized yields in the context of both genetic and management technologies conditional to the frequency of environment types encountered in the TPE. Using ANOVA, it was possible to define domains of application where the opportunity for this approach to close production gaps increase with increasing environmental variability and occurrence of drought stress (Cooper et al., 2020b).

Applications of the CGM-WGP methodology have thus far focused for the most part on field evaluations of a smaller number of populations from maize drought programs (Cooper et al. 2016; Messina et al. 2018). Methodologies that span the scale of the breeding program have been mentioned to be important (Cooper et al. 2016). Complementarily, gap analysis methodologies have been demonstrated in long-term studies for genetic gain but have not been linked to WGP prediction (Cooper et al. 2020b). We propose herein an integrated approach that links the digital tools of CGM-WGP and Gap analysis with MSEs to increase the number of opportunities to realize faster rates of genetic gain in the TPE (Fig 1). Specific objectives of the study were, in the context of a very large breeding half-diallel GxE experiment for maize, to: (1) introduce a strategy for the use of MSEs and MET data in CGM-WGP training and assess CGM-WGP predictive abilities in this context, (2) to introduce the concept of in-silico germplasm characterization, and (3) to connect the Gap analysis and CGM-WGP methodologies to create a tool through which breeders could select the best combinations of genotype and management to close productivity gaps in the TPE.

## Results

### Training strategy: Simulating average performance of genotypes and environment types in MSE and MET

Complex systems modeling can generate mathematical artifacts. A first evaluation of the underlying CGM consists of using parameters known for commercial hybrids or the maturity of the breeding population to check for simulation accuracy across environment types. Baseline simulations of yield (Y_s_) in each environment (Table 1) approximated mean yield of the population (Y_o_) within 15% error (*Y*_*o*_ = 93(±122) + 0.95(±0.08) × *Y*_*s*_; *df* = 21; *r*^2^ = 0.86). Using CGM outputs such as the soil water supply that depends on determinants of water balance and root exploration, and the plant demand for water that depends on potential growth and vapor pressure deficit (VPD), it is possible to calculate a daily water supply/demand ratio (S/D) to characterize environmental drought status. Figure 3 shows the daily S/D dynamics for each environment included in the study. Based on the intensity of the stress (reduction of S/D) and the timing relative to the critical developmental period for kernel set determination in maize, it was plausible to identify three water deficit (WD) environments with low S/D values around flowering time at MSE sites, three well-watered (WW) environments at MSE sites, and 17 TPE environments (Figure 3). The WD environments experienced a decrease in water S/D ratio around flowering that was not observed in the WW and TPE environments, except for E9. Therefore, the multienvironment testing in the TPE under sampled drought environments (Fig. 3b).

**Figure 3.**
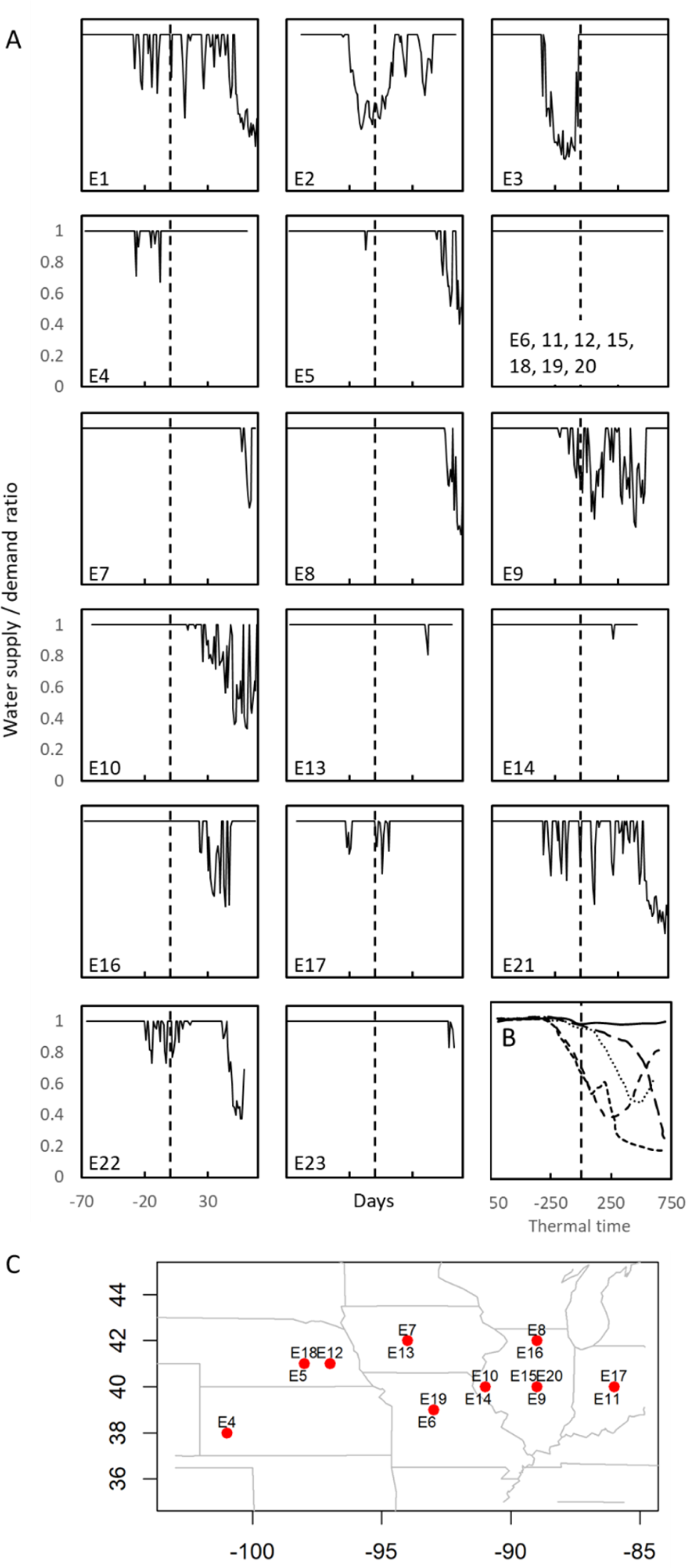
Daily sequences of water supply to demand ratio (S/D, 1=no stress) centered at flowering by environment (E1-E23). Dashed lines indicate flowering. Because S/D was equal to 1 throughout the season E6, 11, 12, 15, 18, 19, 20 were grouped together. Environment types shown in thermal time centered at flowering time (adapted from Cooper et al.,2014b; B). Corn belt testing locations shown in panel C.

**Table 1.**
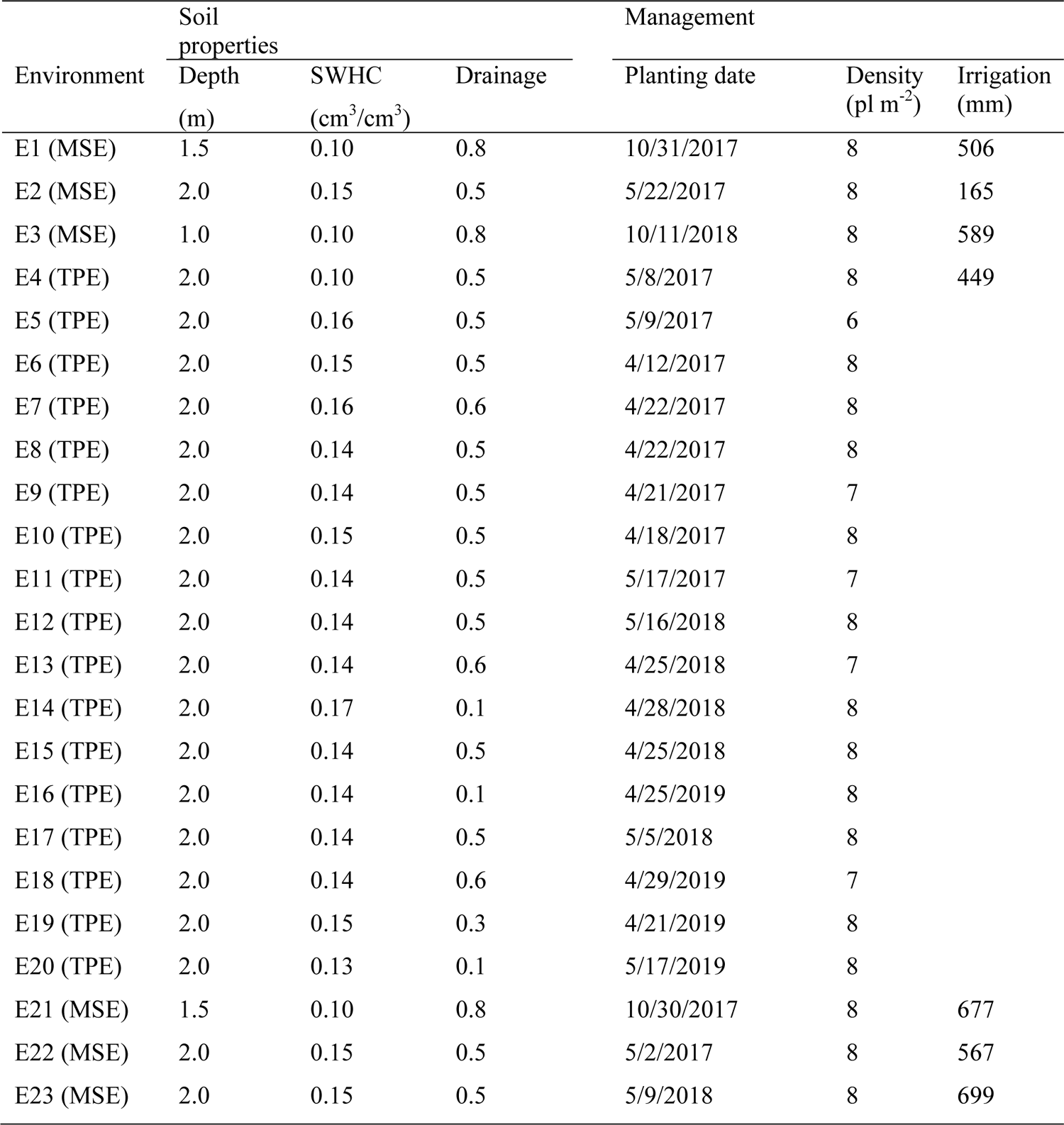
Soil properties and management practices by environment. SWHC: soil water holding capacity

### Training strategy: Harnessing MSE, physiological knowledge, and digital tools to train CGM-WGP

A procedure to make physiological knowledge revealed through CGM-WGP accessible to decision makers and to inform selection decisions is proposed. Similar to forward variable selection in linear regression, a scan of physiological traits is conducted and followed by the combination of these until a parsimonious and physiologically plausible set of traits that minimize the prediction error is identified, and a clear advantage in predictive skill is demonstrated. The selection of candidate traits is informed by observed genotypic variation, as characterized by prior probability distributions. Predictive skill advantage is defined as the difference between the correlation coefficients between observations and predictions for CGM-WGP and WGP at each environment. From the initial scan of 12 model parameters representing key physiological traits, a minimal set was identified that exhibited high correlations between fitted and observed values for yield across environments (Figure 4). This minimum set was comprised of number of kernel rings per ear (NRINGS; high values indicating more kernel rings and sink potential), husk length (HLENGTH; high values indicating long husks), senescence response to water deficit (SENS; low values indicating staygreen), and root elongation rate (RER; high values indicating rapid root elongation). This four-trait CGM-WGP model exhibited correlations between model-generated and observed values that were greater than or at parity with those of WGP (Figure 4); the latter is representative of BayesA. The predictive skill measure clearly demonstrates that for specific population and environment combinations integrating plant physiology into the prediction algorithm contributed to the enhanced modeling of GxE by CGM-WGP in this large experiment.

**Figure 4.**
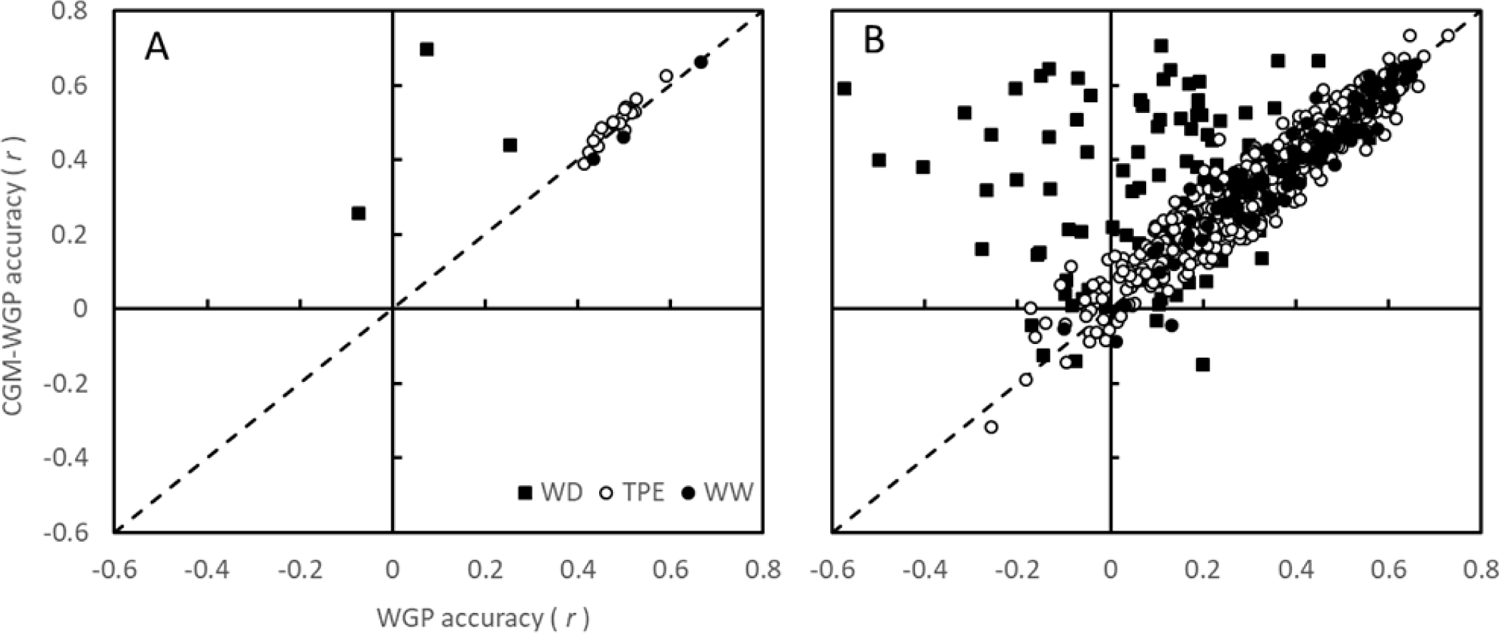
Across family (A) and within family (B) prediction accuracies estimated by the Pearson correlation coefficient (*r*) for whole-genome prediction (WGP) and crop growth model – whole-genome prediction (CGM-WGP) by environment type; water deficit (WD), Target Population of Environments (TPE), and well-watered (WW).

### Training assessment: Predictive skill advantage increased with increasing water deficit

Experimentation in MSEs brings the opportunity to improve phenotyping by eliciting targeted physiological responses in the germplasm subject to selection. Expressed trait phenotypes resulting from the differential management realized in the MSEs (Fig. 3a) enables the estimation of CGM parameters that encapsulates the mechanisms that underpin germplasm performance in the TPE, which include environments with various types of water deficit (Fig. 3b). Goodness-of-fit and predictive ability advantage of the four-trait CGM-WGP model relative to WGP was highest when both tested genotypes and environments were included as part of the subset of data used for training the algorithms (Table 2). Goodness-of-fit and predictive ability advantage decreased, similarly for any combination of untested genotypes, environments and their combination. The difference in the correlation coefficients (*r*_CGM-WGP_ – *r*_WGP_) varied between 0.17 and 0.19 when training the model using data from both water deficit and irrigated experiments at MSEs (Table 2). Excluding water deficit data from the training data set increased goodness-of-fit and predictive ability of WGP (Table 2) because of the contrasting genetic correlations (*r*_G_) between irrigated and TPE (*r*_G_=0.68) and water deficit and TPE experiments (*r*_G_=0.0003), and the under sampling of water deficit environments in the TPE (Fig. 3a).

**Table 2.** Average predictive skill of crop growth model wholegenome prediction (CGM-WGP) and wholegenome prediction (WGP) for cases resulting from the combination of: tested (T) and untested (U; not included in the training of the model) genotypes and tested and untested environments

| Environment | Well-watered data |  |  |  | Well-watered & water deficit data |  |  |  |
| --- | --- | --- | --- | --- | --- | --- | --- | --- |
|  | CGM-WGP |  | WGP |  | CGM-WGP |  | WGP |  |
|  | Genotype |  | Genotype |  | Genotype |  | Genotype |  |
|  | T | U | T | U | T | U | T | U |
| T | 0.79 | 0.61 | 0.72 | 0.61 | 0.80 | 0.58 | 0.49 | 0.42 |
| U | 0.42 | 0.42 | 0.41 | 0.41 | 0.43 | 0.42 | 0.24 | 0.25 |

Training the model with data from all 23 environments, within-family predictive abilities were also high with the four-trait CGM-WGP model and tended to be either at parity with those of WGP or greater—with the latter particularly being the case in WD environments (Fig 4b). Taken together, the results show that for this large MET, the predictive skill advantage of CGM-WGP over WGP increased with increasing severity of water deficit (Fig. 5). Whole season total evapotranspiration was used as a measure of the severity of water deficit for each environment. Analyses over 35 families and 23 environments demonstrated that CGM-WGP offered a predictive advantage (y, *r*_*CGM*−*WGP*_ − *r*_*WGP*_) compared to WGP that increased with decreasing evapotranspiration (ET; log(*y*) = 0.80(±0.6) − 0.006(±0.001) × *ET*; *r*^2^ = 0.59; *df* = 21).

**Figure 5.**
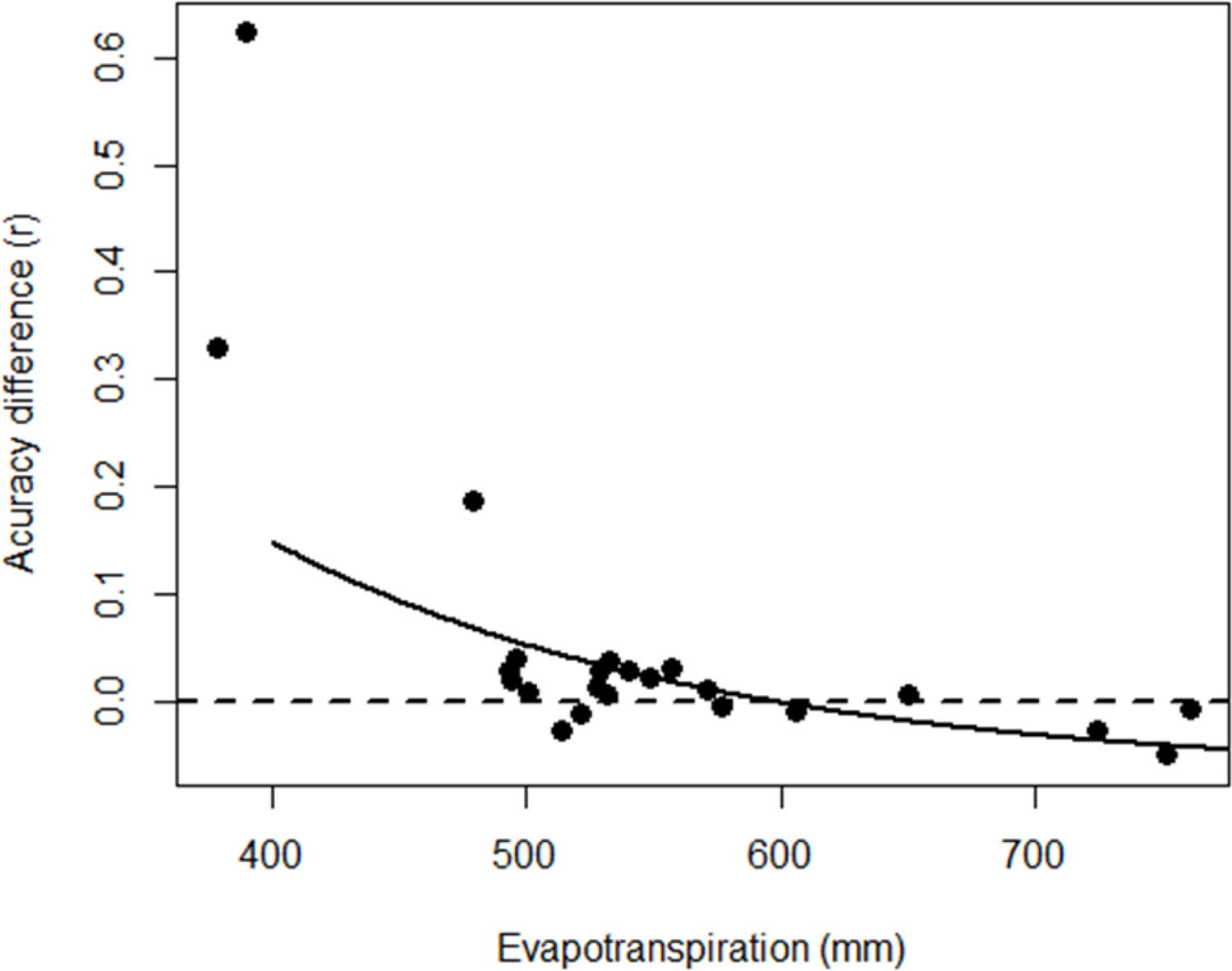
Predictive skill difference in an across-family context estimated by the difference in Pearson correlation coefficients (*r,* observed vs. predicted yield) for each of crop growth model – whole-genome prediction (CGM-WGP) and whole-genome prediction (WGP) methodologies (i.e., *r*_CGM-WGP_ – *r*_WGP_) as a function of evapotranspiration. Each point represents the mean difference in prediction accuracy for across families in a single environment. All environments were included for training and prediction

### Training assessment: Predictive skill in CGM-WGP correlate with robustness of trait estimate

Two of the four estimated traits, NRINGS and SENS, were further examined for the purpose of determining extent of trait stability, when varying the environment types included in the training set and the number of traits being estimated simultaneously. Both of these methodological points are important in the training and application of CGM-WGP and Gap methodology. When including all of the data generated in the MSE environments (WW + WD) in the training set, the posterior ranges for the mean of NRINGS and SENS corresponded closely to their respective prior ranges for the mean. When estimating only one of these traits at a time, use of all MSE environments (WW + WD) vs. only WW environments in the training set produced similar NRINGS estimates (*r*=0.99) but vastly different SENS estimates (*r*=0.26). In the opposite scenario, use of all MSE environments (WW + WD) vs. only WD environments in the training set produced similar SENS estimates (*r*∼1.0) but vastly different NRINGS estimates (*r*=0.49), when estimating only one of these traits at a time. A similar pattern in stability of trait estimates within and across environment types was observed when estimating both NRINGS and SENS simultaneously. To examine from the perspective of number of estimated traits when including only WW environments in the training set, NRINGS estimates were stable whether estimating only NRINGS or both NRINGS and SENS (*r*=0.99). The same was found for WD environments and the stability of SENS estimates, whether estimating only SENS or both NRINGS and SENS (*r*=0.99). When using only the TPE environments as the training set, estimates of NRINGS in the 2017 vs. 2018 TPE environments were moderately to highly stable (*r*=0.90).

### In silico germplasm characterization: relating genetic with functional diversity as determinants of yield

Estimated physiological traits were weakly correlated with each other in pairwise examinations (Table 3) and principal component analysis (PCA) biplots (Fig. 6). NRINGS was positively correlated with SENS and HLENGTH, indicating that a stronger sink was associated with higher senescence and longer husks. RER was positively correlated with SENS, suggesting rapid root elongation was associated with reduced staygreen, and negatively correlated with HLENGTH, suggesting rapid root elongation was associated with improved synchrony of silk exertion and pollen release in WD. Yield under WW conditions in both MSEs and the TPE was strongly correlated with NRINGS (Fig. 3A, Table 3), a determinant of sink potential in the CGM, but not under WD. Yield under WD in MSE was strongly correlated with SENS when severe stress occurred prior to flowering or grain filling, indicating a limitation in source (Fig. 3A, E1 and E3, Table 3). In contrast, SENS and yield in the TPE were positively correlated, suggesting that SENS captured the remobilization due to the establishment of a strong sink rather than a source limitation. When WD occurred around flowering time (Fig. 3A, E2), yield under WD was negatively associated with HLENGTH due to the relationship between silk elongation rate under water deficit and the distance required for silk exposure to pollen (HLENGTH). Timing of water deficit in the MSE around silking (Fig. 3, E2) likely exposed genetic variation in husk length affecting the timing and synchrony of pollination akin to the negative relationship between anthesis-silking interval and grain yield.

**Figure 6.**
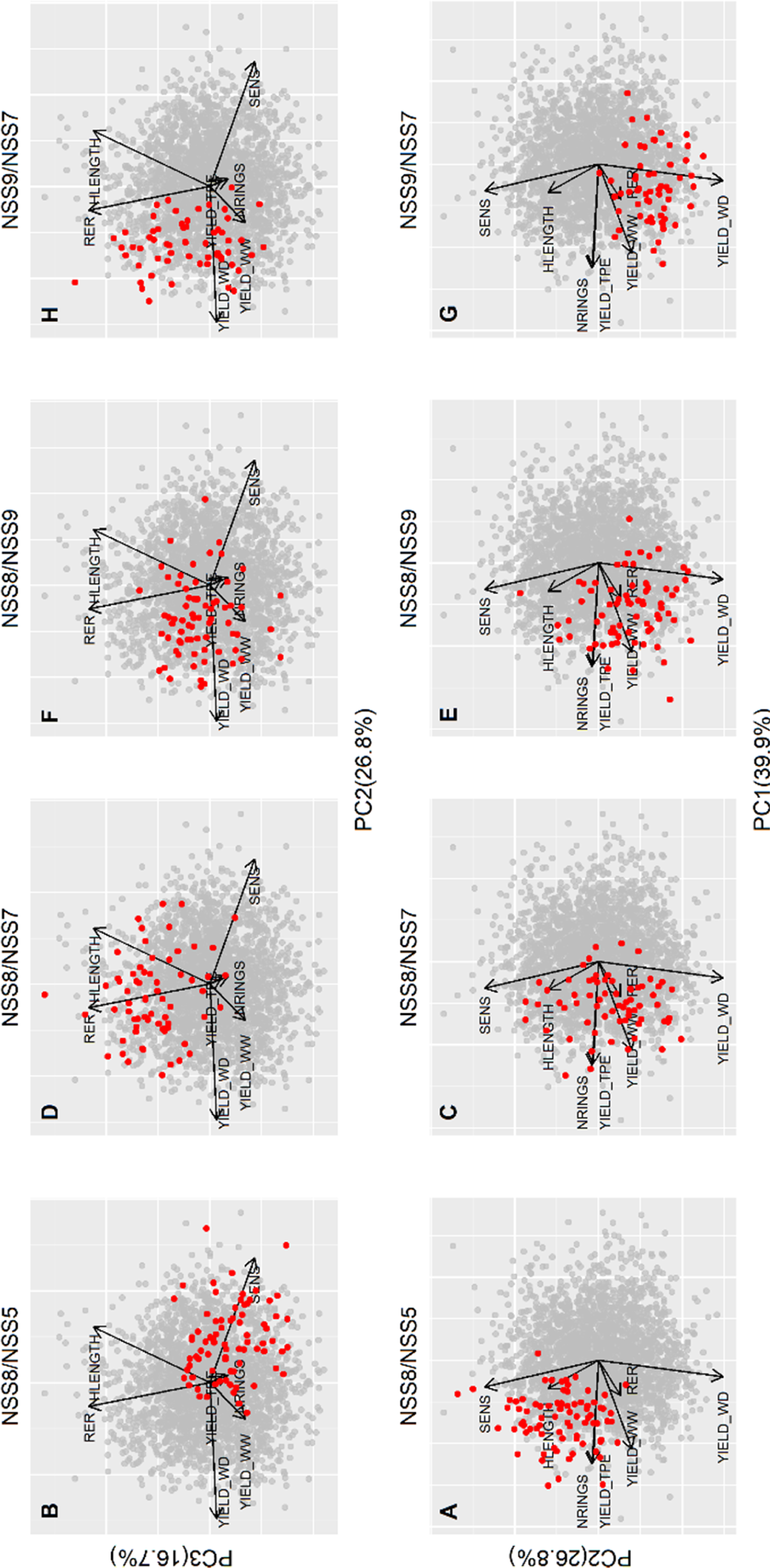
Biplots for principal components (PC) 1, 2 and 3. All hybrids included in the study are represented by grey dots. Selected crosses shown as red dots and visualized in different panels. Vectors are for yield under well-watered conditions, water deficit at flowering time (WD), and in the target population of environments (TPE), and for estimated physiological traits: number of rings per ear (NRINGS), leaf senescence response to water deficit (SENS; negative values indicate staygreen), root elongation rate (RER) and length of husk (HLENGTH).

**Table 3.** Genetic variance (upper triangle) and Pearson correlation coefficients (lower grid; all pairwise combinations) for estimated physiological traits number of rings per ear (NRINGS), leaf senescence response to water deficit (SENS), root elongation rate (RER) and length of husk (HLENGTH), and yield under water deficit, well-watered conditions, and in the multienvironment trial (MET) conducted in the target population of environments (TPE)

|  |  | NRINGS | SENS | RER | HLENGTH |
| --- | --- | --- | --- | --- | --- |
| NRINGS |  | 10.6 | 0.24 | 0.10 | 0.28 |
| SENS | | | $5.1 \times 10^{-5}$ | 0.32 | -0.07 |
| RER |  |  |  | 0.25 | -0.27 |
| HLENGTH |  |  |  |  | 56.6 |
| Yield water deficit | Chile | -0.05 | -0.80 | -0.07 | 0.06 |
|  | USA | -0.02 | -0.05 | 0.04 | -0.66 |
| Yield well-watered | USA & Chile | 0.72 | -0.07 | -0.03 | 0.11 |
| Yield MET | USA TPE | 0.96 | 0.20 | 0.03 | 0.35 |

PCA biplots were used to visualize the relationship among traits with yield in four contrasting populations resulting from crossing high yielding (NSS8 and NSS5) and drought tolerant parents (NSS7 and NSS9) in three environment types (Fig. 6). The first, second and third components explained 39.9%, 26.8% and 16.7% of total G+GxE variance, respectively. PCA1 discriminated hybrids for yield under WW, TPE and NRINGS. PCA2 discriminated hybrids for yield under WD and SENS. PCA3 discriminated hybrids for RER and HLENGTH. Most hybrids from the cross NSS8/NSS5 were high yielding under TPE and WW but low yielding under WD (Fig. 6a,b). Most hybrids had low scores for RER and SENS, with SENS and yield under WD being negatively correlated (Fig. 6b). Crossing NSS8 with drought tolerant parent NSS7 generated a population of hybrids with high scores for yield in the TPE, about 50% of hybrids with high scores for yield under WD (Fig. 6c), RER and HLENGTH, and most hybrids having low scores for SENS, which translates in maintenance of a green canopy under water deficit (Fig. 6d). The population NSS8/NSS9, another cross of NSS8 with a drought tolerant parent, produced a high frequency of hybrids with high scores for yield in both the TPE and WD (Fig. 6e), and consistently low scores of SENS (Fig. 6f). In comparing NSS8/NSS9 with the other crosses with NSS8, the scores for HLENGTH were not changed relative to NSS8/NSS5 but decreased relative to NSS8/NSS7 (Fig. 6b,f). Hybrids resulting from crossing two drought tolerant parents (NSS9/NSS7) had consistently high scores for yield under WD (Fig.6g) and RER (Fig. 6h), and low scores for SENS and HLENGTH (Fig. 6h).

Overall, different traits contribute to germplasm adaptation to water deficit and well-watered conditions in MSE and the TPE, and the germplasm sampled in this study exposed genetic variation for these traits. The examples presented showed the possibility to improve yield under WD by improving simultaneously at least two traits related to capture of water (RER), maintenance of the canopy (SENS) and synchronous timely pollinations, in this case expressed by low HLENGTH. It appears as well that improvement for yield potential via NRINGS could indeed incur benefits for yield int the TPE. Adaptation and expression of GxE for yield emerge from different physiological pathways.

### Integrated framework: Gap analysis and CGM-WGP methodologies can help breeders close the productivity gap

Grain yields in the TPE and the two well-watered environments were all near the 80% quantile front used to define the realistic bound for efficient production agriculture. Average evapotranspiration (ET) was between 492 and 649 mm in the TPE environments, while it increased to 700-800 mm in the MSE well-watered environments (Fig 7a). This sample of environments is highly biased when considering the types and frequency of environments expected in the TPE (Fig. 3; Cooper et al., 2020b). Deviations of the average yields relative to the 80% quantile front indicated gaps in yield productivity across all environments but were more evident under low ET (Fig. 7a). Cross-over GxE interactions for yield performance were observed across ET levels among the families. For example, the NSS8/NSS5 family had low mean yield at low ET levels (806 g m^-2^ for ET < 480 mm) and higher mean yield at high ET levels (1635 g m^-2^) relative to other crosses between NSS8 with drought tolerant parents; yield at low and high ET levels were in the range 892-978 g m^-2^ and 1575-1607 g m^-2^, respectively (Fig 7). Under low ET the yield advantage (291 g m^-2^) of crosses with drought tolerant parents relative to NSS8/NSS5 was highest in E3 (Fig. 3) when severe water deficit occurred preflowering. In contrast, because water deficit occurred around flowering time in E2 (Fig. 3), yield advantage varied among crosses: −68, 3 and 14 g m^-2^ for NSS8/NSS7, NSS8/NSS9 and NSS9/NSS7, respectively. The full expression of drought tolerance related to high RER scores in NSS8/NSS7 (Fig. 6d) did not occur until it was combined with a reduced HLENGTH (Fig. 6h) that enabled a timely pollination (NSS9/NSS7) and with consistent low SENS values (Fig 6g). Note that the yield advantage for NSS9/NSS7 was greater than those of the NSS8/NSS7 and NSS8/NSS9 crosses (Fig. 6). This result demonstrates the opportunity to be purposeful about closing productivity gaps with respect to limited natural/production resources by means of crop improvement.

**Figure 7.**
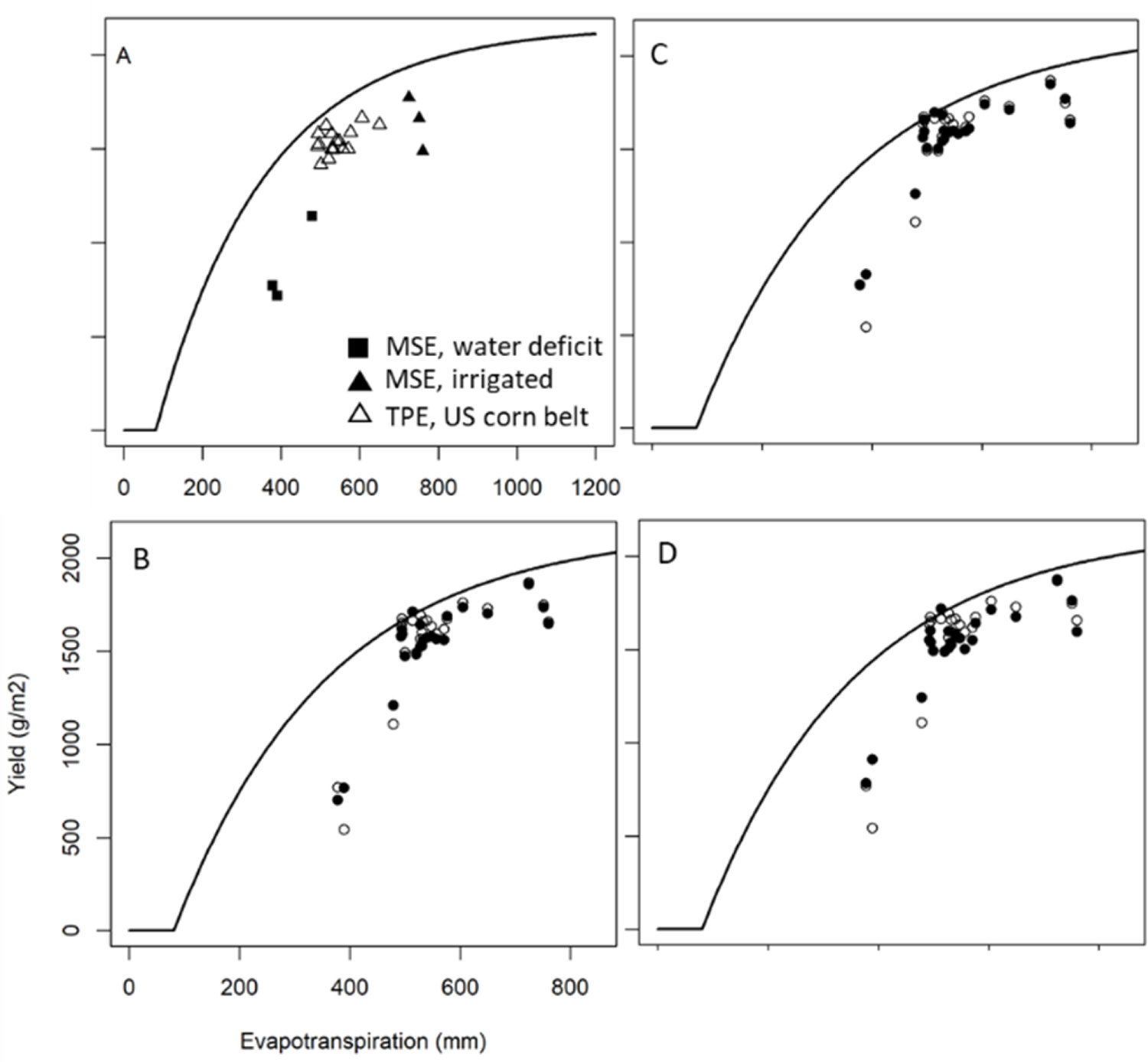
Gap analyses relative to the 80th percentile yield front demonstrated for the average yields across families at each environment (A, Cooper et al., 2020b), and for four contrasting crosses along an evapotranspiration gradient: NSS8/NSS7 (B), NSS8/NSS9 (C), NSS9/NSS7 (D), and NSS8/NSS5 (B-D; open symbol).

Because hybrids were characterized genetically and physiologically, it is possible to use the predictors from CGM-WGP to simulate the performance of each hybrid under different managements to identify opportunities to further close the production gap (Fig 1).

## Discussion

Here we demonstrated an integrated approach that links digital and field experimental approaches using in combination CGM-WGP, Gap analysis, and MSEs to hasten genetic gain (Fig 1). Using a large dataset comprising of 23 locations that exposed 2367 maize hybrids to a range of water deficit and well-watered environments, we estimated that the average out-of-sample predictive skill, both genotype and environment, for WGP and CGM-WGP were 0.25 and 0.42, respectively (Table 2). Here we provide empirical evidence for the robustness of predictive ability of CGM-WGP with changing environments, in contrast with WGP, that is consistent with results from simulation (Messina et al., 2018). Considering the genomic breeder’s equation as a valid framework to quantify the value of the information and the prediction approach (Voss-Fels et al., 2019), the gain in predictive skill due to the use of physiological knowledge to model GxE translates into an average differential response to selection 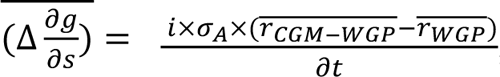, where 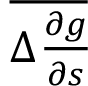 is the average differential genetic gain per unit cycle of selection, *i* is the standardized selection differential, *σ*_*A*_ is the square root of the additive genetic variance in the training population, and *r*_*k*_ are the average correlations between the predicted yields for method *k* and the corresponding values in the TPE. A positive difference in the correlations implies a gain in skill due to modeling main effects and GxE interactions. Because for the germplasm used in this study the gain in predictive skill increased with increasing water deficit (Fig. 5), the gains are dependent on the frequency of environment types and magnitude of GxE. Considering these and prior results (Cooper et al., 2016; Messina et al., 2018) we propose that with access to a suitable CGM, linking genomics and physiology should lead to 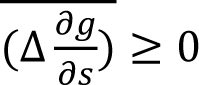. The introduction of a method for model selection, akin to forward variable selection in statistics, enables practitioners other than physiologists to apply CGM-WGP in breeding programs thus increasing the opportunities to expand the application of the method to other germplasm and crops and geographies. The CGM-WGP framework enables the integration of phenomic data that will contribute to overcoming limitations to translate advanced phenomics into genetic gain (Araus et al., 2018). The use of MSE to generate appropriate environmental conditions to elicit physiological responses is a core requirement to generate stable parameters for the selected model and the associated estimation of allelic effects, and to generate environments that expose productivity gaps to inform selection decisions. Over three years of experimentation (Fig 7a) most of the MET results sampled well-watered environmental conditions conducive to high yields (Fig 3a,b). While these experiments are useful for selection, ignoring the biased sampling of the TPE is conducive to missing opportunities to accelerate genetic gain either due to underestimating G, GxE, and prediction accuracy of methods such as CGM-WGP. The integration of Gap analysis with a simulation step, using allelic effects estimated for each G, and E and M intensively sampled from the TPE (e.g., 10^8^; Cooper et al., 2020b), enables implementation of a weighted selection methodology to account for the sampling bias, as advocated by Podlich et al. (1999). Current computing capabilities should be adequate to implement digital phenotyping as proposed here on millions of G, E and M combinations for any crop for which only genetic information on relatedness to tested genotypes information is available. Finally, the CGM-WGP approach can assist in starting answering questions regarding the adaptation of any G or GxM combination to current and/or future climates and production systems, which is not possible using conventional empirical sampling approaches but requires connecting genomics and physiology as demonstrated here.

### Strategy for the use of MSEs and MET data in CGM-WGP training for prediction

This study tested the empirical application of CGM-WGP in a large maize breeding population, with yield as the observed emergent property to be used in model training. The approach is generalizable to the use of a combination of complex traits such as grain yield across diverse environments, moderately complex traits such as leaf area, and directly measured constants for simple traits such as parameters for light response curves (Fig 2). Runs of the CGM, parameterized for checks or more generally the germplasm of interest, were conducted to examine baseline yield simulation and confirm the reasonableness of environmental inputs. Estimation of model parameter vectors in the G, E, and M scenarios of interest were key to implementation of the predictions for physiological traits segregating in the breeding populations. The feasibility of identifying a minimum parsimonious set of parameters for genetic modeling greatly facilitates the routine application of the approach when compared with previous efforts (Cooper et al., 2016; Messina et al., 2018). The use of MSE provided critical information to improve the estimation of CGM parameters, in agreement with prior results (Messina et al., 2018).

Estimation of yield in METs can also help with trait parameterization and model identifiability (Technow et al. 2015), which could be particularly pertinent in instances where influences of emergent properties other than yield cannot be observed, e.g. the influence of enhanced rate of silk elongation and kernel set in low ET drought affected environments contained within the MET.

With the incorporation of WGP and a sampling component, attributes of plant growth that were not measured in the field but that were influential to yield can be appropriately approximated in genotype-specific fashion by integrating over time the rates of growth, development and biomass partitioning. This sampling procedure can be trained on yield alone, with few additional field measurements needed for the CGM-WGP to be used. The refinement of priors for physiological traits can be conducted on a small subsample of individuals that are representative of the breeding population. The parameterization of variation for physiological traits through CGM-WGP is contrasted with more intensive approaches in which all individuals in a breeding population are directly phenotyped for the physiological traits of interest (Yin et al., 2000; Reymond et al. 2003, Messina et al., 2006, Chenu et al. 2009, Messina et al. 2011). The CGM-WGP framework can thus be used to introduce, —following initial experimentation to refine priors,—a physiological component into the analysis of field trials in one or multiple stages of breeding programs (Fig. 1), where in a typical season it may only be economical and/or logistically feasible to quantify yield for the majority of the tested genotypes. Further, advancements in high throughput phenotyping for canopy (Rutkoski et al., 2016; Crain et al., 2017), photosynthesis (Yendrek et al., 2017; Cotrozzi et al., 2020), reproductive (Gage et al., 2017; Berghoefer et al., 2020), and quality (Tillman et al., 2006) traits can create opportunities for a hybrid approach between direct phenotyping at the field level (Messina et al., 2011) and digital characterization of germplasm as demonstrated in the present study (Fig. 6). This integration can address what was recognized as a problem of translating phenomics into decisions in breeding (Araus et al., 2018). We hypothesize that such an approach can increase predictive skill by reducing the underspecification of data-driven models and facilitating a deeper understanding of the physiological determinants of adaptation in the germplasm, and genetic determinants of physiological processes.

### Evaluating CGM-WGP accuracy using a very large breeding half-diallel GxE experiment

The WGP approach used herein represents a reasonable benchmark for CGM-WGP, in that it reflects contemporary, purely statistical methods for prediction of yield from marker effects (Meuwissen et al. 2001; Lorenz et al. 2011; Voss-Fels et al. 2019) as they are applied in commercial maize breeding (Cooper et al. 2014b). The results presented in this study are applied to a significantly larger set of G and E scenarios for the TPE than previous studies that used MSE data for at most four populations (Cooper et al., 2016; Messina et al., 2018). For an experiment comprising 23 locations and 2367 maize hybrids, representing 35 populations, we demonstrated a decrease in the accuracy difference between BayesA, a widely used method, and CGM-WGP with decreasing water deficit (Fig. 5). However, in agreement with previous studies (Cooper et al., 2016; Messina et al., 2018), whenever GxE was small, CGM-WGP still performed at parity with linear models (Fig. 4; Table 2). Therefore, in agreement with a previous study (Messina et al., 2018), it was possible to obtain a robust estimation of physiological traits with the use of multiple environment types, ranging from drought to well-watered, in the training set. Together, these findings suggest that CGM-WGP offers utility in incorporating signals from multiple environment types, and that the difference between benchmark methods and a form of CGM-WGP (Messina et al., 2018; Millet et al., 2019; van Eeuwijk et al., 2019) will increase with the increasing importance of GxE, GxM, and GxExM interactions in the determination of yield.

### inSilico germplasm characterization for hasten genetic gain

Understanding physiology at the level of individual genotypes offers utility both in germplasm characterization and in making selections that maintain physiological diversity, for risk management in the short, medium, and long term (Hammer et al. 2020). However, physiological experiments often focus on few genotypes, due to the intensiveness of the phenotyping methods and/or the systems-level of detail that is required to build a comprehensive mechanistic understanding. Linking genomics and physiology through CGM-WGP brings opportunities to generate hypotheses about the mechanisms of adaptation for millions of untested individuals for which only marker information is available (Fig. 1). Here we focused on four traits for estimation using the CGM-WGP model: NRINGS, HLENGTH, SENS, and RER, and estimated values for 2367 hybrids at 23 locations (Fig 6). A principal component analysis showed that estimated parameters using the CGM-WGP are physiologically sound, and exposed genotypic variation within the germplasm (Fig 6, Table 4).

Because NRINGS affects the potential number of silks, it is an important trait in defining yield potential (Messina et al., 2019). Results conform to the expected positive relationship between NRINGS and yield when water deficit is low (Fig 3; Fig 6). Because silks must extend beyond the husk, for a given elongation rate husk length can determine protandry, failure in pollination and low yields (Hall et al., 1982; Messina et al. 2019) as exposed by the trait relationships depicted in PCA biplots (Fig 6).

HLENGTH and tightness can also determine susceptibility to ear diseases, which was not considered in the model. This trait could thus be somewhat informative for differential yield performance in TPE environments, and a weak positive correlation was indeed observed (Fig. 6). SENS models the response of leaf area loss to water S/D, and contributes to maintenance of photosynthetic rates, grain growth and yield under certain water deficit environments (Borrell et al., 2000; Duvick et al., 2005; Messina et al., 2020a). RER can impact the timing and volume of soil water available to the plant (Hammer et al. 2009, Lynch 2013). Since the introduction of the hypothesis that deep roots contribute to long-term genetic gain for yield of maize in the US corn-belt, two studies (Reyes et al., 2015; Messina et al. 2020a) have shown that total water extraction itself was not found to have changed over 50 years of maize breeding despite substantial genetic gain for yield, such that other mechanisms of yield optimization were likely exploited by breeding (Reyes et al. 2015; Messina et al., 2020a).

Because RER varied among populations (Table 2), water deficit was imposed during the critical window for yield determination in the Woodland MSE (Fig. 3a, E2) and the depth of water extraction in Woodland can occur to depths greater than 2.5 m (Table 1, Reyes et al., 2015), results from this experiment allowed testing of the hypothesis that RER is correlated with yield under WD in deep soils. On average, the yield difference between NSS8/NSS7 and NSS8/NSS5, which have contrasting scores for yield under WD and RER, was negative (−67 g m^-2^) and consistent with prior results (Reyes et al., 2015). However, the yield advantage due in part to high scores in RER (Fig. 6d) was not fully expressed but except in a population (NSS9/NSS7) with low scores for HLENGTH and SENS (Fig. 6g,h). HLENGTH and other traits are determinants of a timely pollination (Messina et al., 2019) contributing to reproductive resilience. These results suggest the hypothesis that in the absence of limitations to root growth in the soil profile (Ordóñez et al. 2018, Osborne et al. 2020, Fan et al. 2016), and considering that reproductive resilience underpins long-term genetic gain (Messina et al., 2020a), the maintenance of gains in reproductive resilience will hasten genetic gain for yield when combined with positive selection for RER.

### On the future of Gap analysis and CGM-WGP digital methodologies

The CGM-WGP framework unifies the extent of physiological detail developed regarding crop growth and development on a daily timescale, with the germplasm testing and selection strategies that already take place within plant breeding pipelines (Technow et al. 2015, Messina et al. 2011). Here we extended the system to consider the Gap between attained and potential yield for a given availability of a yield limiting resource, in this case water. We demonstrated that Gap analysis was useful in examining levels of yield performance in the various environments analyzed in this study, and in identifying families that tended to display more or less stability in yield performance across water availability levels measured by crop ET. These findings related to yield performance and stability can also be examined in light of the characterizations provided by the CGM-WGP model. For example, certain combinations of parents may tend to alleviate or exacerbate one or more trait vulnerabilities (e.g. for NSS8/NSS5, in the case of SENS) or bolster or weaken certain strengths. Continued integration of gap analysis methodologies with CGM-WGP could thus provide insight into specific targets for improvement of yield and yield stability across environmental gradients in the TPE. Considering the farming system context can provide a productive next methodological step to realize crop improvement gains through changes in root systems (Thorup-Kristensen and Kirkegaard, 2016; Bančič et al., 2021), which so far have been elusive in maize (Reyes et al., 2015; Messina et al., 2020a).

Robust predictive abilities of the CGM-WGP methodology were observed both across and within families (Fig. 4), and predictive abilities and physiological trait estimates were stable upon inclusion of data from multiple environment types in the model training data set. The outputs of the CGM-WGP framework additionally enabled germplasm characterization and gap analysis, which provided insight into opportunities for further improvement of yield and yield stability through breeding and/or agronomy.

These findings suggest the multi-faceted utility of CGM-WGP in large breeding populations and early stages of the breeding process and later stages of product placement (Fig. 1) for the continued improvement—with potential increases in efficiency and genetic gain, as is enabled by predictive skill— of yield and yield stability in the TPE.

## Conclusion

Based on the results from the analyses of a very large maize dataset we conclude that the integration of physiological understanding improves predictive skill for the TPE. The advantage of the CGM-WGP approach increased when water deficit environments were involved and decreased with decreasing water deficit, or with more generally decreasing contributions of GxE, GxM, or GxExM to the total variance. Integrated systems approaches can facilitate the application of physiological knowledge in breeding via CGM-WGP. Because plants and breeding systems are evolving complex systems and yield, and other phenotypes of interest are emergent phenotypes of those systems, ongoing research is needed to increase relevant understanding of the physiological basis of adaptation. We have combined physiology and breeding through the CGM-WGP methodology and demonstrated the emerging opportunity to leverage more digital technologies for digital phenotyping for characterization and prediction of germplasm, and to dedicate more resources to advance the scientific understanding of the links between genomics and physiology through modeling.

## Materials and Methods

### Data

A maize breeding and genetics experiment was conducted by crossing nine non-stiff stalk (NSS) inbred parents, denoted as NSS1, NSS2, …, NSS9, in a half-diallel mating design. The resulting 35 families, each of which included about 75 doubled haploids (DHs), were crossed to a common stiff stalk inbred tester resulting in 2367 hybrids in total. These hybrids were evaluated in 23 environments (herein E1 through E23) from 2017 to 2019. Six environments were from managed stress environments (MSE) located at Corteva research stations in Woodland, CA and Viluco, Chile. Planting density, planting date and crop husbandry followed best local practices (Table 1). Planting spacing was 0.76 m for all environments. Irrigation was managed to impose water stress at different times of development including flowering (E1-3; Fig. 3). Well-watered (WW) irrigated controls were included. Irrigation was applied using drip tape buried at 20 cm deep (E21-23, Fig. 3; Table 1). The remaining 17 environments were in the US corn belt states. Yield was measured at each location using mechanical combines and adjusted to 150 g kg^-1^ grain moisture.

Soil data required to run simulations using the CGM were from in-field measurements. Daily weather data (solar radiation, maximum and minimum temperature, and precipitation) were from nearby weather stations from the National Oceanic and Atmospheric Administration (Bell et al., 2013; Table 1). Environment and management parameters such as plant population (plants per square meter), and planting date were also included (Table 1).

### Experimental design and statistical analyses

The experimental design in each environment was a row-column design with diagonal checks. The grain yield data were analyzed using the ASREML mixed model software (Gilmour et al. 2009) for each environment with genotype as a fixed effect, row/column as random effects and AR1xAR1 residual structure,

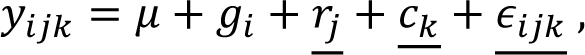

where *y*_*ijk*_ is the yield for genotype *i* in row *j* and column *k*, μ is the overall mean, *g*_*i*_ is effect of genotype *i*, *r*_*j*_ is the effect of row *j*, *c*_*k*_ is the effect of column *k* and, *∈*_*ijk*_ are the residual effects. μ and *g*_*i*_ are fixed effects, *r*_*j*_ and *c*_*k*_ are assumed to be randomly normally distributed variables with mean 0 while *∈*_*ijk*_ are assumed to be randomly normally distributed variables with mean 0 and variance matrix *R* = *σ^2^_e_[AR1(ρ_r_) ⊕ AR1(ρ_c_)]*, representing the Kronecker product of first-order autoregressive processes across rows and columns, respectively, with the spatial residual variance *σ*^2^_e_. Best linear unbiased estimators (BLUEs) for each genotype and for each environment were produced after adjusting for spatial effects and were used for subsequent analyses. Linear regressions were conducting using R (R Core Team, 2020). Principal component analyses were used to characterize physiological parameters estimated for the 35 families and 2367 hybrids and yields under different environment types. Analyses were conducted using the *prcomp* function in R package stats (R Core Team, 2020).

### Genotyping

Each DH was genotyped with approximately 2935 single-nucleotide polymorphism (SNP) markers. Missing SNP allele calls were imputed based on parent-progeny relationship and founder allele frequency.

### CGM-WGP configurations

This research used the CGM-WGP methodology described by Messina et al. (2018). Descriptions for the CGM were reported (Messina et al., 2015; Cooper et al., 2016). An algorithm was included in the CGM to simulate silk elongation and response to S/D ratio. The algorithm is based on cohorting floret/kernel rings in the ear, and pollination after silk emergence from tip of the husk based on pollen availability (Oury et al., 2016; Turc et al., 2016; Messina et al., 2019). Briefly, the maximum number of silks is determined by the number of rings per ear (NRINGS) and kernels per ring (KRINGS). Silk emergence depends on the average rate of silk elongation, its response to water deficit, and the average distance that the average silks needs to travel along the husk (HLENGTH). The availability of pollen at any time follows a Gaussian distribution centered shortly after the time of anthesis/shedding of the main culm (Uribelarrea et al. 2002). Changes with ages in silk receptivity follows Anderson et al., 2004. The simulated total number of embryos determined the attainable harvest index as described in Cooper et al. (2016). The maximum daily leaf senescence fraction response to water deficit is set to 0.05 and decreases with increasing S/D up to zero when S/D equals 1.

The Bayesian hierarchical model was used to model allele effects for physiological traits as described in Messina et al. (2018) with the extension to model soil properties such as depth, which is estimated independently for each location but held constant for all genotypes evaluated at a given location. The rationale for modeling soil factors as a variable is that for many environments sampled in a plant breeding trial these model inputs are either unknown or known with under-desired precision.

Inaccurate environmental inputs directly impact the accuracy of crop growth model simulation and thus prevent the accurate estimation of physiological traits and the genetic determinants. Allowing important environmental inputs to be estimated jointly with physiological traits prevents the model from exploring unrealistic physiological trait space because of biased environmental inputs. Prior distributions were from Messina et al. (2018). The prior for soil depth was a truncated normal distribution with 0 m as the lower bound and 2000 mm as the upper bound for each location, and variance of 25 cm. The Metropolis-Hastings-within-Gibbs algorithm was used to sample all parameters, including soil depth.

### Model evaluation and selection

Models, which herein refer to a CGM-WGP models for which different CGM parameters were estimated using marker data, were evaluated for their capacity to describe and predict observed trait phenotypes. The metric to compare simulations and observations was the Pearson correlation coefficient. Evaluations were conducted for the prediction of mean yield performance for all genotypes across locations, for the prediction of family means across environment types (water deficit, corn belt trials/TPE, irrigated), and for the prediction of hybrids across environment types. Comparisons between WGP and CGM-WGP were conducted to assess advantages of using MSEs and CGM-WGP in prediction. Eight cases (Table 2) stem from the prediction of family means for the observations in the corn belt (or TPE) using data collected in MSEs (irrigated, and irrigated plus water deficit data): the combination of tested environments and genotypes (included in the training of CGM-WGP or WGP), untested (or out-of-sample; not included during the training process) environments and genotypes. Genotypes used to train the model were a random sample of 250 out of the total of 2467 hybrids.

A trait selection scheme was designed to identify feasible sets of physiological traits that have acceptable predictive skill. This method is deemed necessary both biologically and computationally. Biologically, not all twelve traits are relevant in the present data set. If all environments were well-watered, traits related to yield potential should be more informative whereas if all environment were water-limited, traits related to drought tolerance should be more informative. In the case of sets of environments comprised of contrasting water conditions, both yield potential traits and drought tolerance traits should be needed to understand and model the GxE variation. Moreover, traits impacting different physiological processes in the CGM may result in similar yield variation observed and thus may have similar importance. Including multiple or all traits may result in model unidentifiability issues given that different combinations of traits could tend to result in similar model likelihoods. The trait selection scheme can be viewed as a nonlinear analog of forward variable selection in multiple linear regression. Since this model evaluation procedure involved many runs of CGM-WGP, only 250 randomly chosen genotypes were used in the trait selection procedure to reduce run time. This procedure starts with only one trait in the model. To evaluate the fitness of each of the twelve one-trait models, by-location accuracy was calculated as the correlation between the predicted yield and observed yield and compared with BayesA by-location accuracy where the BayesA WGP model (Meuwissen et al. 2001) was applied directly to the by-location BLUEs for yield averaged over all locations. The results can be inspected in scatterplots with BayesA by-location accuracy on the x-axis and the one-trait model by-location accuracies on the y-axis. Upon inspection and calculation of prediction accuracies, the most predictive one-trait model is selected. Other traits are selected in an iterative manner for their potential to increase the model goodness of fit. A limited number of candidate multi-trait models were evaluated, and one parsimonious set was selected for the purpose of comparing CGM-WGP with BayesA results.

Twelve CGM parameters coding for rates regulating physiological processes were tested as the candidate traits driving the yield variation and GxE: 1) leaf appearance rate (Muchow et al. 1990; based on analysis of data from Messina et al., (2011); μ = 0.00275, *σ*^2^ = 1.62 10^−8^; leaf °C^-1^), 2) thermal time using base temperature 0 for grain fill duration (based on Gambín et al. (2006); μ = 1300, *σ*^2^ = 2603; °C), 3) radiation use efficiency (based on Messina et al. (2018); μ = 1.85, *σ*^2^ = 0.16; g MJ^-1^), 4) area of the largest leaf in the canopy (based on Messina et al. (2018); μ = 850, *σ*^2^ = 650; cm^2^), 5) number of rings per ear (NRINGS; based on analysis of data from Messina et al. (2011) assuming 16 kernels per ring; μ = 45, *σ*^2^ = 6.5), 6) slope above breakpoint describing the relative (0-1 where 1=no response) transpiration response to VPD (informed by results from Choudhary et al. (2014) expressed on relative scale; μ = 0.5, *σ*^2^ = 0.0026), 7) breakpoint of transpiration response to VPD (informed by Messina et al. (2015); μ = 2, *σ*^2^ = 0.065; kPa), 8) senescence coefficient (SENS) which reduces leaf area in a linear manner in accordance with the water S/D (μ = 0.05, *σ*^2^ = 0.0001; dimensionless), 9) maximum silk elongation rate per hour (informed by results from Turc et al. (2016); μ = 1.5, *σ*^2^ = 0.065 cm h^-1^), 10) fraction of total soil water when silk elongation rate was reduced to 50% of maximum (informed by results from Turc et al. (2016); μ = 0.5, *σ*^2^ = 0.01; dimensionless), 11) husk length (HLENGTH; informed by Messina et al. (2019); μ = 200, *σ*^2^ = 104; mm), 12) root elongation rate (RER, informed by data from Dardanelli et al. (1997), Hammer et al. (2009), Singh et al. (2010), van Oosterom et al. (2016) and Ordóñez et al. (2018); μ = 25, *σ*^2^ = 6.5; mm d^-1^).

## Data

The data can be made available through https://openinnovation.corteva.com/ upon reasonable request for public research purposes and project evaluation.

## Acknowledgements

Sandra Truong for simulating maize canopy growth and development used to create the panel A in Figure 2. To the many colleagues who planted, sprayed, and harvested the field trials. John Arbuckle, Geoff Graham and Matt Smalley supported the project.

## Literature cited

1. Anderson SR, Lauer MJ, Schoper JB, Shibles RM (2004) Pollination timing effects on kernel set and silk receptivity in four maize hybrids. Crop Sci 44:464–473

2. Araus JL, Kefauver SC, Zaman-Allah M, Olsen MS, Cairns JE (2018) Translating high-throughput phenotyping into genetic gain. Trends Plant Sci 23:451–466

3. Bančič J, Werner CR., Gaynor RC, Gorjanc G, Odeny DA., Ojulong HF., Dawson IK, Hoad SP, Hickey JM (2021). Modeling illustrates that genomic selection provides new opportunities for intercrop breeding. Front. Plant Sci. doi: 0.3389/fpls.2021.605172

4. Bell, JE, Palecki MA, Baker CB, Collins WG, Lawrimore JH, Leeper RD, Hall ME, Kochendorfer J, Meyers TP, Wilson T, et al (2013) U.S. Climate Reference Network soil moisture and temperature observations. J Hydrometeorol 14:977–988

5. Berghoefer, CC, Hanselman TA, Hausmann NJ, Messina C (2020) Methods of yield assessment with crop photometry. US Patent 10,713,768

6. Chapman SC, Cooper M, Butler D, Henzell R (2000) Genotype by environment interactions affecting grain sorghum. I. Characteristics that confound interpretation of hybrid yield. Aust J Agric Res 51:197–208

7. Chenu K, Chapman SC, Tardieu F, McLean G, Welcker C, Hammer GL (2009) Simulating the yield impacts of organ-level quantitative trait loci associated with drought response in maize: a “gene-to-phenotype” modeling approach. Genetics 183:1507–23

8. Choudhary S, Sinclair TR, Messina CD, Cooper M (2014) Hydraulic conductance in maize hybrids differing in breakpoint of transpiration response to increasing vapor pressure deficit. Crop Sci 54: 1147–1152

9. Comstock RE (1977) Quantitative genetics and the design of breeding programs. In E Pollak et al., ed, Proceedings of the international conference on quantitative genetics. Iowa State University Press, USA, Iowa, pp 705–718

10. Cooper M, Woodruff DR, Eisemann RL, Brennan PS, DeLacy IH (1995) A selection strategy to accommodate genotype-by-environment interaction for grain yield of wheat: managed-environments for selection among genotypes. Theoret Appl Genetics 90, 492–502

11. Cooper M, Gho C, Leafgren R, Tang T, Messina C (2014a) Breeding drought-tolerant maize hybrids for the US corn-belt: discovery to product. J Exp Bot 65:6191–6204

12. Cooper M, Messina CD, Podlich D, Totir LR, Baumgarten A, Hausmann NJ, Wright D, Graham G (2014b) Predicting the future of plant breeding: complementing empirical evaluation with genetic prediction. Crop Pasture Sci 65:311–336

13. Cooper M, Technow F, Messina C, Gho C, Totir LR (2016) Use of crop growth models with whole-genome prediction: application to a maize multienvironment trial. Crop Sci 56:1–16

14. Cooper M, Powell O, Voss-Fels KP, Messina CD, Gho C, Podlich DW, Technow F, Chapman SC, Beveridge CA, Ortiz-Barrientos D, Hammer GL (2020a) Modelling selection response in plant breeding programs using crop models as mechanistic gene-to-phenotype (CGM-G2P) multi-trait link functions. in silico Plants, diaa016. doi: 10.1093/insilicoplants/diaa016

15. Cooper M, Tang T, Gho C, Hart T, Hammer G, Messina C (2020b). Integrating Genetic Gain and Gap Analysis to predict improvements in crop productivity. Crop Sci 60:582–604

16. Cotrozzi L, Peron R, Tuinstra MR, Mickelbart MV, Couture JJ (2020) Spectral phenotyping of physiological and anatomical leaf traits related with maize water status. Plant Physiol 184:1363–1377

17. Crain J, Reynolds M, Poland J (2017) Utilizing high-throughput phenotypic data for improved phenotypic selection of stress-adaptive traits in wheat. Crop Sci 57: 648–659

18. Dardanelli JL, Bachmeier OA, Sereno R, Gil R, (1997). Rooting depth and soil water extraction patterns of different crops in a silty loam Haplustoll. Field Crops Res., 54: 29–38

19. de la Vega AJ, Chapman SC (2001) Genotype by environment interaction and indirect selection for yield in sunflower: II. Three-mode principal component analysis of oil and biomass yield across environments in Argentina. Field Crops Research 72, 39–50

20. Duvick DN (2005) The contribution of breeding to yield advances in maize (*Zea mays* L.). Adv. Agron. 86, 83–145

21. Fan J, McConkey B, Wang H, Janzen H (2016) Root distribution by depth for temperate agricultural crops. Field Crops Research, 189: 68–74

22. Fischer T, Byerlee D, Edmeades G (2014) Crop yields and global food security: Will yield increase continue to feed the world? ACIAR Monograph No. 158. Australian Centre for International Agricultural Research. Canberra

23. Gambín BL, Borrás L, Otegui ME (2006) Source–sink relations and kernel weight differences in maize temperate hybrids. Field Crops Research 95, 316–326

24. Gage JL, Miller ND, Spalding EP, Kaeppler SM, de Leon N (2017) TIPS: a system for automated image-based phenotyping of maize tassels. Plant Methods 31, 13:2

25. Gilmour AR, Gogel BJ, Cullis BR, Thompson R (2009) ASReml user guide release 3.0 VSN International Ltd, Hemel Hempstead, HP1 1ES, UK.

26. Hall AJ, Vilella F, Trapani N, Chimenti C (1982) The effects of water stress and genotype on the dynamics of pollen-shedding and silking in maize. Field Crops Research 5, 349–363

27. Hammer GL, Dong Z, McLean G, Doherty A, Messina C, Schusler J, Zinselmeier C, Paszkiewicz S, Cooper M (2009) Can changes in canopy and/or root system architecture explain historical maize yield trends in the US Corn Belt? Crop Sci 49:299–312

28. Hammer G, Messina C, Wu A, Cooper M. 2019. Biological reality and parsimony in crop models – why we need both in crop improvement! in silico Plants 1:diz010

29. Hammer GL, McLean G, van Oosterom E, Chapman S, Zheng B, Wu A, Doherty A, Jordan D (2020) Designing crops for adaptation to the drought and high-temperature risks anticipated in future climates. Crop Sci 60:605–621

30. Leakey ADB, Uribelarrea M, Ainsworth EA, Naidu SL, Rogers A, Ort DR, Long SP (2006) Photosynthesis, productivity, and yield of maize are not affected by open-air elevation of CO2 concentration in the absence of drought. Plant Physiol 140:779–790

31. Li X, Guo T, Mu Q, Li X, Yu J. 2018. Genomic and environmental determinants and their interplay underlying phenotypic plasticity. Proc Natl Acad Sci U S A 115:6679–6684

32. Lynch JP (2013) Steep, cheap and deep: an ideotype to optimize water and N acquisition by maize root systems. Ann Bot 112:347–357

33. McCormick RF, Truong SK, Rotundo J, Gaspar AP, Kyle D, van Eeuwijk F, Messina CD (2020) Intercontinental prediction of soybean phenology via hybrid ensemble of knowledge-based and data-driven models biorxiv doi: 10.1101/2020.09.22.30650

34. Messina CD, Jones JW, Boote KJ, Vallejos CE (2006) A Gene-based model to simulate soybean development and yield responses to environment. Crop Sci 46:456–466

35. Messina CD, Sinclair TR, Hammer GL, Curan D, Thompson J, Oler Z, Gho C, Cooper M (2015) Limited-Transpiration trait may increase maize drought tolerance in the US Corn Belt. Agron J 107:1978–1986

36. Messina CD, Podlich D, Dong Z, Samples M, Cooper M (2011) Yield–trait performance landscapes: from theory to application in breeding maize for drought tolerance. J Exp Bot 62:855–868

37. Messina CD, Technow F, Tang T, Totir R, Gho C, Cooper M (2018) Leveraging biological insight and environmental variation to improve phenotypic prediction: Integrating crop growth models (CGM) with whole genome prediction (WGP). Eur J Agron 100:151–162

38. Messina CD, Hammer GL, McLean G, Cooper M, van Oosterom EJ, Tardieu F, Chapman SC, Doherty A, Gho C (2019) On the dynamic determinants of reproductive failure under drought in maize. in silico Plants 1(1) diz003

39. Messina C., Cooper M, McDonald D, Poffenbarger H, Clark R, Salinas A, Fang Y, Gho C, Tang T, Graham G (2020a) Reproductive resilience but not root architecture underpin yield improvement in maize. biorxiv doi: 10.1101/2020.09.30.320937

40. Messina C, Cooper M, Reynolds M, Hammer G (2020b) Crop science: A foundation for advancing predictive agriculture. Crop Sci 60:544–546

41. Millet EJ, Kruijer W, Coupel-Ledru A, Alvarez Prado S, Cabrera-Bosquet L, Lacube S, Charcosset A, Welcker C, van Eeuwijk F, Tardieu F (2019) Genomic prediction of maize yield across European environmental conditions. Nat Genet 51: 952–956

42. Muchow RC, Sinclair TR, Bennett JM (1990) Temperature and solar radiation effects on potential maize yield across locations. Agron J 82: 338–343

43. Mwiinga B, Sibiya J, Kondwakwenda A, Musvosvi C, Chigeza G (2020) Genotype x environment interaction analysis of soybean (Glycine max (L.) Merrill) grain yield across production environments in Southern Africa. Field Crops Res 256:107922

44. National Academies of Sciences, Engineering, and Medicine (2019) Science Breakthroughs to Advance Food and Agricultural Research by 2030. The National Academies Press, Washington, DC: doi: 10.17226/25059.

45. Ordóñez RA, Castellano MJ, Hatfield JL, Helmers MJ, Licht MA, Liebman M, Dietzel R, Martinez-Feria R, Iqbal J, Puntel LA, Córdova SC, Togliatti K, Wright EE, Archontoulis SV (2018) Maize and soybean root front velocity and maximum depth in Iowa, USA. Field Crops Res 215:122–131

46. Osborne SL, Khim Chim B, Riedell WE, Schumacher TE (2020) Root length density of cereal and grain legume crops grown in diverse rotations. Crop Sci doi: 10.1002/csc2.20164

47. Oury V, Tardieu F, Turc O (2016) Ovary apical abortion under water deficit is caused by changes in sequential development of ovaries and in silk growth rate in maize. Plant Physiol 171:986–996

48. Pinheiro J, Bates D, DebRoy S, Sarkar DR (2020) Core Team. nlme: Linear and nonlinear mixed effects models. R package version 3.1–144

49. Podlich DW, Cooper M, Basford KE (1999) Computer simulation of a selection strategy to accommodate genotype-environment interactions in a wheat recurrent selection programme. Plant Breed 118:17–28

50. Poland J, Endelman J, Dawson J, Rutkoski J, Wu S, Manes Y, Dreisigacker S, Crossa J, Sánchez-Villeda H, Sorrells M, Jannink J-L (2012) Genomic selection in wheat breeding using genotyping-by-sequencing. Plant Genome 5:103–113

51. Poland, J (2015) Breeding-assisted genomics. Curr Opin Plant Biol 24:119–124

52. Ramirez-Villegas J, Molero Milan A, Alexandrov N, Asseng S, Challinor AJ, Crossa J, van Eeuwijk F, Edmond M, Grenier GC, Heinemann AB et al (2020) CGIAR modeling approaches for resource-constrained scenarios: I. Accelerating crop breeding for a changing climate. Crop Sci 60:547–567

53. R Core Team (2020) R: A language and environment for statistical computing. R Foundation for statistical computing, Vienna, Austria. https://www.R-project.org

54. Ray DK, Mueller ND, West PC, Foley JA (2013) Yield trends are insufficient to double global crop production by 2050. PLoS One 8: e66428

55. Reyes A, Messina CD, Hammer GL, Liu L, van Oosterom E, Lafitte R, Cooper M (2015) Soil water capture trends over 50 years of single-cross maize (Zea mays L.) breeding in the US corn-belt. J Exp Bot 66:7339–7346

56. Reymond M, Muller B, Leonardi A, Charcosset A, Tardieu F (2003) Combining quantitative trait loci analysis and an ecophysiological model to analyze the genetic variability of the responses of maize leaf growth to temperature and water deficit. Plant Physiol 131:664–675

57. Rincent R, Malosetti M, Ababaei B, Touzy G, Mini A, Bogard M, Martre P, Le Gouis J, van Eeuwijk F (2019) Using crop growth model stress covariates and AMMI decomposition to better predict genotype-by-environment interactions. Theoret Appl Genetics 132: 3399–3411

58. Robert P, Le Gouis J, The BreedWheat Consortium, Rincent R (2020) Combining crop growth modeling with trait-assisted prediction improved the prediction of genotype by environment interactions. Front Plant Sci 11: 827. doi: 10.3389/fpls.2020.00827

59. Rutkoski J, Poland J, Mondal S, Autrique E, Pérez LG, Crossa J, Reynolds M, Singh R (2016) Canopy temperature and vegetation indices from high-throughput phenotyping improve accuracy of pedigree and genomic selection for grain yield in wheat. G3 6:2799–2808

60. Singh V, van Oosterom EJ, Jordan DR, Messina CD, Cooper M, Hammer GL (2010) Morphological and architectural development of root systems in sorghum and maize. Plant Soil 333:287–299

61. Technow F, Messina CD, Totir RL, Cooper M (2015). Integrating crop growth models with whole genome prediction through approximate bayesian computation. PLoS One 10:e0130855

62. Thorup-Kristensen K, Kirkegaard J (2016) Root system-based limits to agricultural productivity and efficiency: the farming systems context. Ann Bot 118:573–592

63. Tillman BL, Gorbet DW, Person G (2006), Predicting oleic and linoleic acid content of single peanut seeds using near-infrared reflectance spectroscopy. Crop Sci 46:2121–2126

64. Turc O, Bouteillé M, Fuad-Hassan A, Welcker C, Tardieu F (2016) The growth of vegetative and reproductive structures (leaves and silks) respond similarly to hydraulic cues in maize. New Phytol 212:377–388

65. Uribelarrea M, Cárcova J, Otegui ME, Westgate ME (2002) Pollen Production, Pollination Dynamics, and Kernel Set in Maize. Crop Sci 42:1910–1918

66. van Eeuwijk FA, Bustos-Korts D, Millet EJ, Boer MP, Kruijer W, Thompson A, Malosetti M, Iwata H, Quiroz R, Kuppe C, Muller O, et al (2019) Modelling strategies for assessing and increasing the effectiveness of new phenotyping techniques in plant breeding. Plant Science 282:23–39

67. van Ittersum MK, Cassman KG, Grassini P, Wolf J, Tittonelli P, Hochman Z (2013) Yield gap analyses with local to global relevance—A Review. Field Crops Res 143:4–17

68. van Oosterom EJ, Yang Z, Zhang F, Deifel KS, Cooper M, Messina CD, Hammer GL (2016) Hybrid variation for root system efficiency in maize: potential links to drought adaptation. Funct Plant Biol 43:502–511

69. Voss-Fels KP, Cooper M, Hayes BJ (2019) Accelerating crop genetic gains with genomic selection. Theoret Appl Genetics 132: 669–686

70. Washburn JD, Burch MB, Valdez Franco JA (2020) Predictive breeding for maize: Making use of molecular phenotypes, machine learning, and physiological crop models. Crop Sci 60: 622–638

71. Wu A, Hammer GL, Doherty A, von Caemmerer S, Farquhar GD (2019) Quantifying impacts of enhancing photosynthesis on crop yield. Nat Plants 5:380–388

72. Yendrek CR, Tomaz T, Montes CM, Cao Y, Morse AM, Brown PJ, McIntyre LM, Leakey ADB, Ainsworth EA (2017) High-Throughput phenotyping of maize leaf physiological and biochemical traits using hyperspectral reflectance. Plant Physiol 173:614–626

73. Yin X, Chasalow S, Dourleijn CJ, Stam P, Kropff MJ (2000) Coupling estimated effects of QTLs for physiological traits to a crop growth model: predicting yield variation among recombinant inbred lines in barley. Heredity 85:539–549

